## Supplementary material for "Intermediaries in phosphotransfer: structural features discriminating hybrid histidine kinase Rec domains from response regulator homologs": Supplemantry Tables (from another version of this preprint: version 1, 2022-06-03)

**Supplementary Table 1. Crystallographic data collection and refinement statistics**

| | |
|---|---|
| Data collection | |
| Synchrotron source | SLS, PXI |
| Wavelength (Å) | 1.00004 |
| Space group | I 1 2 1 |
| a, b, c (Å) | 37.4, 42.9, 65.0 |
| α, β, γ (°) | 90.0, 94.4, 90.0 |
| Resolution (Å) | 28.2-1.25 (1.29-1.25) |
| No. of unique reflections | 28065 (2818) |
| No of reflections used for $R_{free}$ | 1372 (136) |
| Completeness (%) | 98.37 (99.58) |
| I/σ (I) | 17.24 (4.96) |
| Redundancy | 2.0 (2.0) |
| $R_{merge}$ (%) | 1.7 (12.9) |
| $R_{pim}$ (%) | 1.7 (12.9) |
| CC (1/2) (%) | 99.9 (94.0) |
| Refinement | |
| $R_{work}/R_{free}$ (%) | 20.6/22.7 |
| RMSD from ideal values | |
| -- Bond length (Å) | 0.0123 |
| -- Bond angles (°) | 1.7 |
| Molecules/asymmetric unit | 1 |
| No. of atoms | |
| -- Protein | 839 |
| -- Water | 93 |
| Average B-factor (Å$^2$) | 18.9 |
| -- Protein (Å$^2$) | 17.5 |
| -- Water (Å$^2$) | 31.7 |
| Ramachandran statistics | |
| -- Favored regions (%) | 100.0 |
| -- Allowed regions (%) | 0.0 |
| -- Disallowed regions (%) | 0.0 |
| PDB Code | 6TNE |

*) The values recorded in parentheses are those for the highest resolution shell.

**Supplementary Table 2. Amide proton linewidth of $CckA^{Rec}$ residues in the inactive and active state.**

| Residue | Linewidth of amide proton (Hz) | |
|---|---|---|
| | native $CckA^{Rec}$ | $BeF_3^-$ modified $CckA^{Rec}$ |
| L574 | 22.3 | 20.6 |
| V576 | 24.9 | 21.4 |
| V582 | 24.8 | 18.4 |
| V585 | 19.9 | 15.4 |
| G603 | 24.0 | 23.4 |
| I608 | 18.6 | 16.8 |

**Supplementary Table 3. Primers used to obtain the recombinant plasmids used in this study .**

| ID | Name | Sequence (5'>3') | Length | Tm* (°C) | Exp Ta** (°C) |
|---|---|---|---|---|---|
| **1** | Q5SDM_$CckA^{Rec}$_F | CACCATCACCATCACGGTTCTGGTTCTGG CGCCGGCCGCATCCT | 44 | 75.3 | 72.0 |
| **2** | Q5SDM_$CckA^{Rec}$_R | TCAATGGTGATGATGGTGGTGCTACG CCGCCTGCAGCTGCTGCTT | 45 | 76.3 | 72.0 |
| **3** | Q5SDM_$CckA^{Rec}$_N-histag_F | AAGGAGATATACCATATGGGCCATCAC CATCACCATCACGGT | 42 | 68.0 | 72.0 |
| **4** | Q5SDM_$CckA^{Rec}$_ N-histag_R | TCAATGGTGATGATGGTGGTGCTA CGCCGCCTGCAGCTGCTGCTT | 45 | 76.3 | 72.0 |

* Tm = calculated melting temperature ** Exp Ta = annealing temperature used experimentally
